# A KRAS^G12D^-based organoid model for early oncogenic transformation of the mammary epithelium

**DOI:** 10.64898/2026.11.17.766834

**Authors:** Darby J. Jones, Liam E. Rao, Junior J. West

## Abstract

Modeling cancer initiation, progression, and metastasis has greatly impacted our understanding of fundamental cell and molecular mechanisms driving disease and has led to breakthrough treatments that greatly improve patient outcomes. Understanding the earliest stages of tumor formation is particularly important for uncovering the root causes of oncogenic transformation and tumor evolution. It has been historically difficult to model the cellular dynamics that drive the earliest stages of tumor evolution, largely due to difficulties in culturing and manipulating normal primary cells in 3-dimensional environments, triggering oncogenesis, and observing the resulting effects. Recent advances in 3D organoid culture provide a promising avenue to address these challenges. In this study, we have developed a *Kras* based organoid model of early tumorigenesis. In our model, oncogenic transformation was triggered by Cre-mediated expression of oncogenic KRAS^G12D^ in otherwise healthy murine-derived mammary epithelial organoids. While normal organoids displayed controlled organoid growth and morphogenesis, induction of KRAS^G12D^ caused neoplastic organoid growth that occurred with the absence of lumen formation. Gene expression analysis revealed up regulation of several KRAS effectors, and a broad repertoire of genes associated with tumor development and human breast cancer. Induction of oncogenic *Kras* in our model resulted in heterotypic organoids that possessed KRAS^G12D^ + cells growing alongside KRAS^G12D^- cells. In this context, amplification of KRAS^G12D^ + cells occurred at the expense of KRAS^G12D^- cells. Finally we observed that neoplastic growth of KRAS^G12D^ expressing organoids was associated with a suppression of apoptosis. We believe that our organoid model provides a platform that is suitable to examine the earliest stages of tumor development and can be expanded to study several questions related to cancer biology, prevention studies, therapeutic development, and beyond.

**Summary:** In this study, Jones et al., develop a KRAS^G12D^ based organoid model for early stages of oncogenic transformation. Their model shows that early stages of tumorigenesis are associated with induction neoplastic growth, expression of several tumor-associated pathways, including suppression of apoptosis for tumorigenic growth.

## Introduction

Modeling human cancer initiation, progression, and metastasis is critical for defining the cellular mechanisms that drive disease. Various human, animal, and cell culture models of tumor formation, progression, and metastasis have revealed the complex cellular, genetic, and systemic principles that underlie many human cancers [1]. However, studying changes to the cellular landscapes that occur during the earliest stages of oncogenic transformation has been limited to conventional cell culture models that lack the 3-dimensional architectures and cellular heterogeneity that are present in tissues where tumors arise. Organoids have become a powerful model system that can recapitulate the 3D dynamics, cellular interactions, and microenvironmental milieu that models the physiological environment where tumors naturally arise [2, 3]. The modeling capability of organoids also allows for tracking of dynamic cellular processes with high temporal and spatial resolution. Experimental tractability of organoid systems allows for efficient genetic and pharmacological perturbations, co-culture with diverse cell types found in tissues, and physical manipulations, which all contribute to their effectiveness as a model system [4].

Oncogenesis can be triggered by a variety of factors that ultimately lead to genetic aberrations and downstream cellular transformation. Mutations in proto-oncogenes, genes that become activated upon mutation and have the ability to initiate oncogenic transformation, were some of the first genetic aberrations discovered in tumors [5]. Mutant versions of the Ras family of small GTPases were the first oncogenes identified in humans, and KRAS is the most frequently mutated gene in human cancer [6, 7]. Oncogenic mutations are most commonly found in the nucleotide binding p-loop of the protein at amino acid residues G12 and G13. The G12D mutation is the most common and makes up 30% of all KRAS mutations in cancer.

Mechanistically, these mutations limit the intrinsic GTP hydrolysis activity and reduce sensitivity to GTPase activating protein (GAP)-driven GTP hydrolysis, resulting in a constitutively active KRAS [8, 9]. Therefore, mutant KRAS and the resulting effects on downstream signaling pathways is an important area of investigation for cancer biologists, especially in KRAS driven cancers, such as pancreatic ductal adenocarcinoma (PDAC), non-small cell lung cancer (NSCLC), and colorectal cancer (CRC) [7, 10]. Conversely, the role of KRAS in other cancers, such as breast cancer is less well defined. With the recent development of clinically approved RAS inhibitors, it is even more important to understand the role of these proteins in all stages of disease progression [11].

KRAS mutations are only observed in 2%-4% of breast tumors, however, KRAS induced pathways are perturbed in about 20% of breast cancer cases [12–14]. There are a variety of mechanisms that explain why KRAS pathways are up regulated in these tumors in the absence of a specific KRAS mutation. Recently discovered mechanisms include, mutations in downstream RAS effectors (I.e. PI3K), up regulation of upstream receptor tyrosine kinases (i.e. HER2), copy number alterations, down regulation of pathway inhibitors, up regulation of KRAS transcription, and miRNA fluctuations [9, 15, 16]. Therefore, modeling ectopic KRAS pathway activation using oncogenic KRAS is a valuable method for studying the role of KRAS associated pathways in breast cancer initiation, progression, and metastasis. Many models have been developed based on exogenous expression of KRAS with oncogenic point mutations. These models allow for the growth of various breast cancer subtypes *in vivo* [17–19]. However, the majority of these models depend on the epithelial-specific mouse mammary tumor virus (MMTV) promoter [20]. Therefore, any deviation from epithelial identity, for example during epithelial to mesenchymal transitions (EMT), are negatively selected against during tumor progression. The increased invasion and metastasis observed in triple negative breast cancer (TNBC) is associated with increased KRAS pathway activity and EMT [16], making the development of TNBC models based on MMTV-driven expression of KRAS less feasible. Some models do address this discrepancy, where Rädler *et al.* developed an *in vivo* KRAS based mouse model of claudin-low breast cancer that is EMT permissive [21]. While their model is ideal for addressing questions related to *in vivo* tumor initiation, growth, and metastasis, the ability of this model to capture the earlier stages of tumor initiation is limited.

In this study, we developed an organoid model of KRAS driven early oncogenic transformation. This model induces expression of oncogenic KRAS^G12D^ from the endogenous KRAS allele, thereby avoiding the complications of genetic amplifications. By comparing the growth of organoids expressing oncogenic KRAS to control organoids we were able to distinguish features of oncogenic growth from normal growth and morphogenesis of mammary epithelia. We observed that KRAS^G12D^ expression induced neoplastic organoid growth that lacked normal lumen formation within the first 7 days of oncogenic transformation. Gene expression analysis revealed up regulation of a broad range of breast cancer associated genes upon expression of oncogenic KRAS. We also observed heterogeneity in our organoid model where oncogenic KRAS^G12D^ + expressing cells grew alongside normal KRAS^G12D^-cells. Finally, we found that the early stages of oncogenic transformation is associated with a suppression of programmed cell death.

## Results

### Organoids expressing KRAS^G12D^ display increased growth and lack of lumen formation

To investigate how the earliest stages of oncogenic transofmation differ from normal epithelial growth and morphogenesis, we isolated organoids from mice harboring a lox-stop-lox cassette upstream of the *Kras^G12D^* transgene and an mTmG (membrane-Tomato/membrane-GFP) CRE-reporter trans gene (Fig. 1A-B). We incubated these organoids with empty vector adenovirus (Ad-Null) or adenovirus encoding Cre recombinase (Ad-CRE) overnight, plated them in 3D matrigel domes the following day, and cultured the organoids for at least 7 days. We noticed that the media used to culture Ad-CRE treated organoids became orange/yellow within ∼4 days of culture, indicating a pH change due to nutrient consumption, whereas the media used to culture Ad-Null treated organoids remained pink in color (Fig. 1C). We also observed that organoid pellets obtained from the same number of Ad-Null and Ad-CRE organoids after 7 days in culture were noticeably different in size, where Ad-CRE organoid pellets were noticeably larger. These observations suggested that over the course of our organoid assay, the Ad-CRE treated organoids grew at a significantly faster rate.

**Figure 1:**
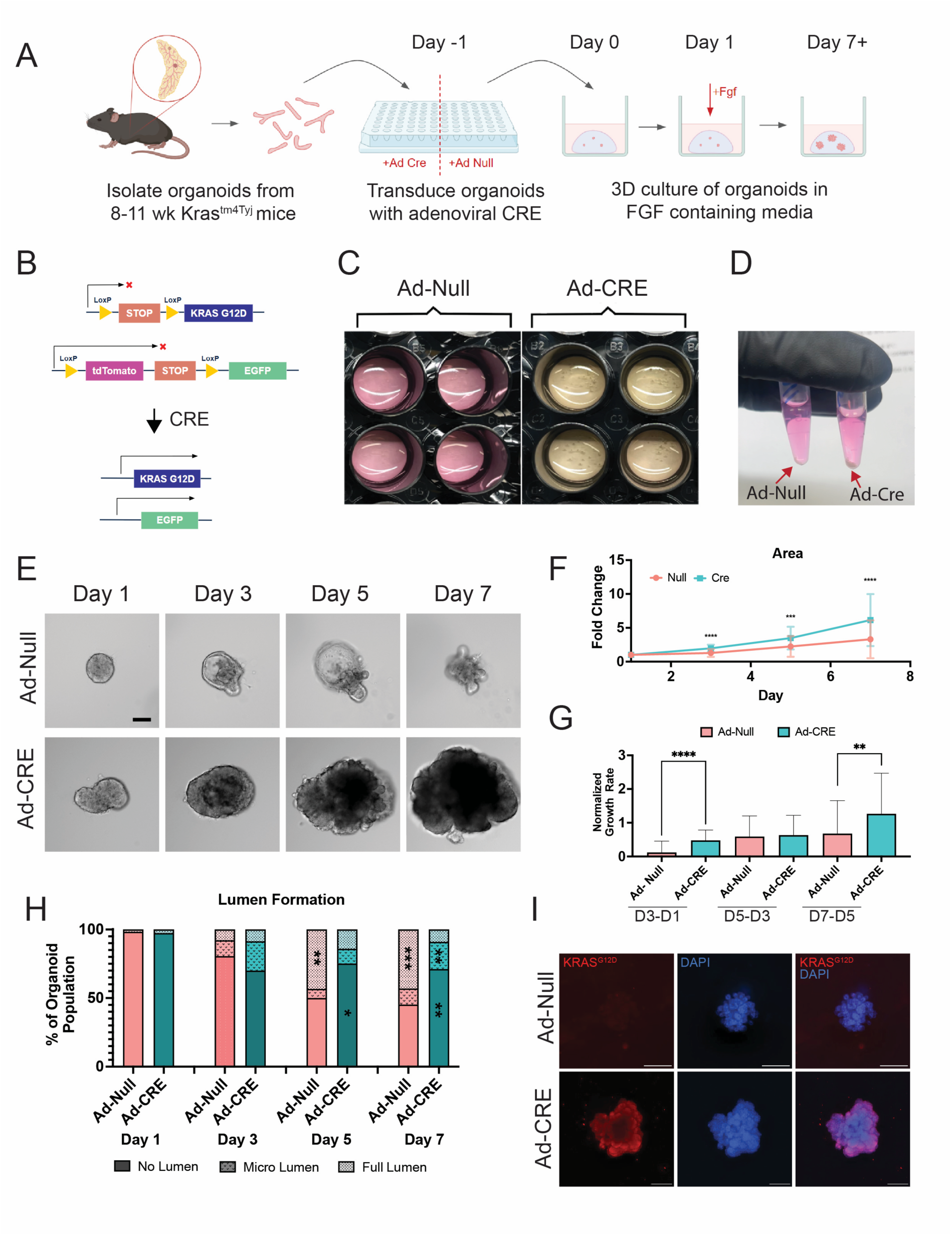
Activation of KRASG12D induces neoplastic organoid growth. (A) Schematic of workflow. (B) Schematic of Cre recombinase inducible KRASG12D mutation (top) and mTmG cre-recombinase reporter (bottom) before and after Ad-CRE transduction. (C) Representative image of wells with media on day 7 of culture for Ad-Null treated organoids (left) and Ad-CRE treated organoids (right). (D) Representative image of pellets of Ad-Null treated (left) and Ad-CRE treated (right) organoids once isolated from Matrigel dome on day 7 of culture. (E) Representative brightfield images of organoids on day 1, day 3, day 5, and day 7 of 3D culture. Scale bar is 100 µm. (F) Quantification of organoid size over time. Each point represents the mean normalized area of organoids from 3 biological replicates (n = 43-48 per condition). Error bars are standard deviation (SD). Mann-Whiteny test was used to determine significance. ***P≤0.001, ****P≤0.0001. (G) Quantification of normalized organoid growth rate over time. r = 3, n = 43-48 per condition. Error bars are SD. Mann-Whiteny test was used to determine significance. **P≤0.01, ****P≤0.0001. (H) Percent of organoids with no lumen, one or more micro lumens, or one or more full lumens on each day. r = 3, n = 46-48 organoids per condition. Chi squared test was performed to determine significance. Significance is indicated on the significantly larger bar. *P≤0.05, **P≤0.01, ***P≤0.001. (I) Representative immunofluorescent images of fixed organoids on day 7. Red is KRASG12D, blue is DAPI. Scale bar is 100 µm.

To directly measure the growth rates of organoids from both conditions, we captured images of organoids on every other day of 3D culture (Fig. 1E). By measuring organoid area over time, we observed a significant increase in organoid size over time (Fig. 1F) and average organoid growth rate (Fig. 1G), where Ad-CRE organoids grew faster than Ad-Null organoids overall, with specific increases between days 1-3 and days 5-7. In addition to organoid size, we observed differences in organoid morphology over time. While both Ad-Null and Ad-CRE treated organoids looked similar in morphology on day 1, by day 3 about half of the Ad-null treated organoids began to clear out the core of the organoid and generate lumen, resulting in organoids with cystic morphologies by day 5 (Fig. 1E, Supp Fig. 1A). After day 3, we observed the formation of epithelial buds that go on to form elongated branches as has been previously described [22]. In our Ad-CRE treated organoids, although we observed about 10%-20% of organoids forming micro lumen, we rarely observed (<10%) organoids forming full lumen and becoming cystic organoids (Fig. 1E, H). Furthermore, Ad-CRE treated organoids formed fewer elongated branches, resulting in more amorphous organoid shapes. To validate that the effects we observed in organoid growth and morphology were due to expression of Kras^G12D^, we performed immunofluorsence using an antibody that recognizes KRAS^G12D^ specifically. We observed increased fluorescence intensity in Ad-CRE treated organoids when compared to Ad-Null conditions (Fig. 1I).

### KRAS^G12D^ expression induces oncogenic pathways within the first 7 days of induction

Transcriptional targets of KRAS^G12D^ have been well characterized in many systems [9]. Therefore, to determine the transcriptomic response of the mammary epithelium to KRAS^G12D^ expression, we performed bulk RNA-seq on organoids after 7 days of 3D culture. PCA analysis revealed a clear separation between Ad-Null and Ad-Cre treated samples, where the majority of samples clustered with each other based on their experimental condition. This indicates a distinct transcriptomic response of mammary organoids in response to KRAS^G12D^ expression (Fig. 2A). By performing differential gene expression analysis on Ad-Null vs. Ad-CRE samples, we observed a total of 657 significantly differentially expressed genes (Fig. 2B). Next, we applied a gene set enrichment analysis (GSEA) to the RNA-seq data to determine changes to co-regulated gene sets that are induced in response to KRAS^G12D^ expression (Fig. 2C). We observed a significant enrichment in many gene sets that have been previously associated with active Kras, such as ‘Myc targets’, ‘MTORC1 signaling’, ‘KRAS signaling up’, ‘E2F targets’, ‘TNFA signaling via NFKβ’, ’epithelial to mesenchymal transition’, and ‘PI3K AKT MTOR signaling’.

**Figure 2:**
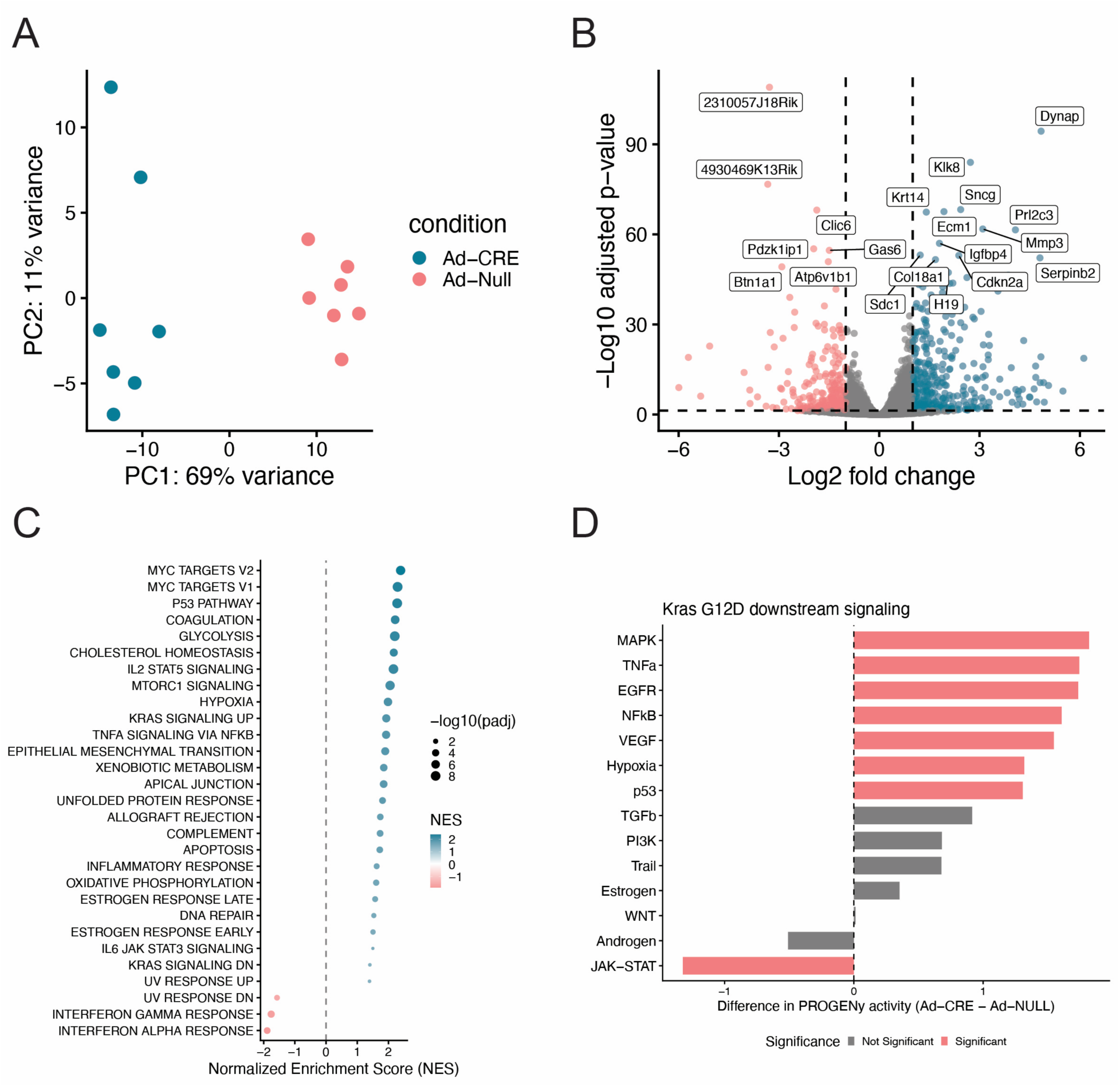
KRASG12D inudction triggers oncogenic signaling and a broader tumorigenic gene expression profile within the first 7 days of organoid culture. (A) Principal component analysis (PCA) plot of Ad-Null versus Ad-CRE total RNA sequences using 3’ end counting via plasmidsaurus. Blue is Ad-CRE, pink is Ad-null. (B) Volcano plot of differentiall expressed genes when comparing Ad-CRE treated organoids to Ad-Null treated organoids. Significance was determined by an adjusted p value <0.05 and a log_2_foldchange < 1. (C) Gene set enrichment analysis (GSEA) of the RNA-seq data. (D) Predicted intracellular pathways upregulated based on PROGENy analysis of sequenced RNA.

To further investigate the intracellular signaling pathways that were activated in response to KRAS^G12D^ expression, we performed a PROGENy analysis, which estimates intracellular signaling pathway activity based on expression of downstream transcriptional targets [23]. The top five up regulated signaling pathways determined by PROGENy were ‘MAPK’, ‘TNF⍺’, ‘EGFR’, ‘NF**κ**B,, and ‘VEGF’, which are all known to act either directly or indirectly up and downstream of Kras signaling [24]. We additionally observed up regulation of ‘Hypoxia’, and ‘p53’ signaling, which have both been implicated in KRAS intracellular signaling [24].

Interestingly, PROGENy did not detect a significant up regulation of ‘PI3K’ signaling, which is also known to act downstream of active KRAS. Therefore, transcriptional response of oncogenic Kras expression in the mammary epithelium primarily relies on the known receptor tyrosine kinase (RTK), MAPK, and TNF⍺/NF**κ**B intracellular signaling pathways associated with Kras, and not others, such as PI3K associated pathways.

Further investigation into the previously ascribed functions of the differentially expressed genes revealed a number of relevant biological characteristics that were induced in KRAS^G12D^ expressing organoids (Supp. Fig. 1C). Notably, we observed a number of genes that have been specifically associated with breast cancer, basal-like/squamous epithelial differentiation, immune involvement, tumor suppression, and invasion/extracellular matrix remodeling.

Therefore, KRAS^G12D^ expression is sufficient to induce widespread gene expression changes that affect many aspects of tumor initiation and progression within the first 7 days of oncogenic transformation.

### Organoid growth occurs by overcoming programmed cell death

To further investigate the cellular dynamics of organoid growth and morphology, we imaged the mTomato-mGFP (mTmG) CRE reporter. As expected, by day 7 of organoid culture, we observed no mGFP expressing cells in the Ad-Null treated organoids, whereas Ad-CRE treated organoids exhibited a high number of mGFP expressing cells (Fig. 3A). We also observed a number of mTomato expressing cells in the Ad-CRE condition, which likely represent cells that were not transduced with Ad-CRE. Upon closer examination of the Ad-CRE treated organoids, we observed that the mTomato expressing took on branched morphologies, whereas the mGFP expressing cells failed to generate budding structures, and were responsible for the overall amorphous shape of the Ad-CRE treated organoids (Fig. 3A). To understand the emergence of these morphologies, we imaged organoids over the course of 7 days (Fig. 3B).

**Figure 3:**
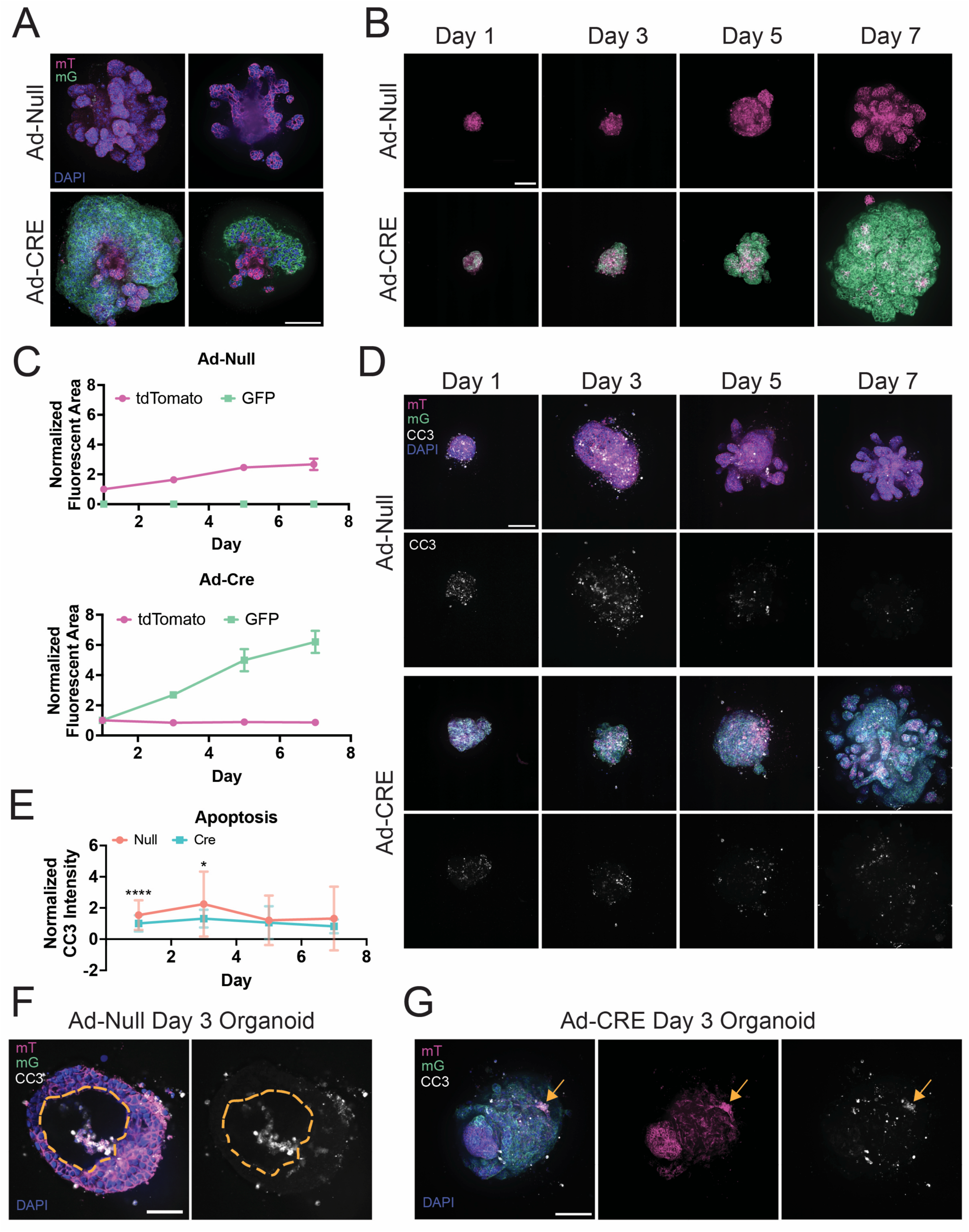
Suppression of apoptosis drives neoplastic growth of heterotypic KRASG12D organoids. (A) Representative fluorescent images of Ad-Null (top) and Ad-CRE (bottom) organoids fixed on day 7. Full z-projections are left and individual z-slices are right. Scale bar is 100 µm. (B) Representative fluorescent images of single organoids images over the course of 7 days of culture. Z-projections are displayed with Ad-Null condition top and Ad-CRE condition bottom. Scale bar is 100 µm. (C) Quantification of mT and mG cell populations over time. Error bars are standard error of the mean (SEM). (D) Representative z-stack projections of organoids fixed and stained with cleaved caspase 3 (CC3) over time. Scale bar is 100 µm. (E) Quantification of CC3 fluorescence over time. Raw integrated density of CC3 within organoid z-projection area was measured and normalized to DAPI raw integrated density. Error bars are SD. Mann-Whiteny test was used to determine significance. *P≤0.05, ****P≤0.0001. (F) Z-slice of panel D day 3 Ad-Null organoid as representative image of CC3-associated lumen formation. Lumen is outlined in the yellow dotted line on both images. Scale bar is 100 µm. (G) Representative image of Ad-Cre organoid on day 3 of culture with co-localization of mT and CC3 signals (indicated by the yellow arrow). Scale bar is 100 µm.

Throughout the time course, Ad-Null treated organoids only contain mTomato expressing cells. However, as early as Day 1, we observe the presence of mGFP expressing cells together with mTomato expressing cells in Ad-CRE treated organoids. As time progresses we observe a significant increase in the number of mGFP expressing cells, with a concomitant decrease in the number of mTomato expressing cells when normalized to day 1 of 3D culture (Fig. 3B-C).

This result indicates that the increased growth of KRAS^G12D^ expressing organoids is due to the expansion of a population of KRAS^G12D^ expressing cells at the expense of cells that lack KRAS^G12D^ expression.

Organoid growth can be explained by increased cell proliferation, decreased cell death, or a combination of the two. To distinguish between these possibilities, we performed immunofluorescence for phospho-histone H3 (pHH3) as a marker for mitotic cells, and cleaved caspase 3 (CC3) as a marker for programmed cell death through the apoptotic pathway. We did not observe a significant difference in the amount of cell division between Ad-Null and Ad-CRE treated organoids on any day of culture (Supp. Fig. 1B). However, we did observe a significant increase in the amount of CC3 signal in the Ad-Null treated organoids on days 1 and 3 of 3D culture (Fig. 3D), indicating that expression of Kras^G12D^ suppresses apoptosis during organoid growth and morphogenesis. We looked closer at the temporal and spatial distributions of apoptotic cells during organoid growth. Interestingly, we observed that in Ad- Null organoids, apoptosis occurs most frequently in cells that are inside of a forming lumen (Fig. 3F). The CC3 signal decreases significantly by Day 5 of 3D culture, which corresponds to when the full lumen/cystic organoid phenotype emerges (Fig. 1E,G). This suggests that lumen formation and cyst morphology is driven by apoptosis. In addition to the decreased CC3 signal observed in Ad-Cre treated organoids generally, we also observed that the CC3 signal that is present is found largely in the mTomato expressing cells that lack KRAS^G12D^ expression (Fig. 3G), further demonstrating that KRAS^G12D^ expression suppresses apoptosis in this context.

## Discussion

In this study we developed a KRAS^G12D^ based organoid model of early oncogenesis. Activation of oncogenic KRAS^G12D^ expression resulted in increased neoplastic growth of mammary epithelial organoids, which included loss of lumen formation within the first 7 days of 3D organotypic culture. RNA-seq analysis of day 7 organoids revealed hundreds of differentially expressed genes between KRAS^G12D^ expressing organoids and control organoids, indicating a robust gene expression signature that differentiates early oncogenic growth, from normal growth and morphogenesis of mammary epithelia. Pathway analyses revealed that this gene expression signature included significant increases in intracellular signaling pathways that are directly and indirectly related to KRAS signaling. Activation of KRAS^G12D^ expression also induced expression of a broad repertoire of tumor associated genes, some specific to human breast tumors, at early stages of oncogenic transformation. We also observed that CRE-treated organoids in our model possess cells that are KRAS^G12D^ positive and negative. At early timepoints, we found that KRAS^G12D^ expressing cells compose the majority of the organoid and are responsible for driving the neoplastic organoid growth. We found that the difference in organoid growth is associated with decreased apoptosis in KRAS^G12D^ expressing cells, where non-expressing cells displayed increased apoptosis whether they were cultured alone or in close contact with KRAS^G12D^ expressing cells.

The relationship between oncogenic transformation and apoptosis is complex and highly context dependent [25]. One of the main features of oncogenic transformation is increased cell survival and decreased cell death [26]. However, in some studies, apoptosis has been shown to be required for tumor formation, and could provide more physical space for breast cancer cells to become more invasive [27]. FAK-induced oncogenic transformation of mammary epithelial cells have also been shown to work through suppression of apoptosis [28]. Although we observed that increased apoptosis was associated with normal mammary epithelial cells, we did observe lower levels of apoptosis in KRAS^G12D^ expressing cells, which does not preclude the importance of apoptosis for oncogenic transformation. The highest levels of apoptosis were associated with lumen formation in our organoid model. Apoptosis-induced lumen formation is a normal feature of mammary epithelial morphogenesis and is an important mechanism to maintain normal epithelial architecture and function [29]. Our observation that expression of KRAS^G12D^ is sufficient to block this process raises the question whether blocking apoptosis-induced lumen formation is a strategy used by cancer cells more broadly. Cell growth in the lumen of mammary ducts is the defining feature of ductal-carcinoma *in situ* (DCIS), the pathological classification of early breast cancer detection. Since our model of early tumorigenesis is likely recapitulating features of DCIS, it may be a suitable to study the onset of this process in more detail. In our gene expression data, we observed up regulation of AKT and IL-6 signaling, which are both anti-apoptotic pathways [30], and may trigger the onset of DCIS.

Myc is an oncogene and master regulator of cell growth, metabolism, and biosynthesis that acts downstream of KRAS. Myc is up regulated in 30-50% of high grade breast tumors, many of these being TNBC and drug resistant tumors [31]. We observed a strong up regulation of many Myc target genes and Myc-associated tumor characteristics in our organoid model, suggesting that transformation is likely occurring through this pathway. Myc can induce stem-cell features [32, 33], E2F expression [34], and EMT [35], all of which are observed in our gene expression analyses. Although we did not observe increased expression of Myc genes themselves, the RAS-MAPK pathway promotes Myc signaling by phosphorylation of Ser62, which promotes its stabilization and allowing it to exert its downstream effects. Therefore, of the many downstream KRAS effectors, Myc signaling is likely a critical route to transformation in our organoid system.

Our current proposed model relies on 3D culture of isolated mammary epithelial organoids. Tumor initiation occurs in a tissue context that includes several non-epithelial tissue components, also referred to as the tumor microenvironment that are not included in our model. However, we observed significant changes in the expression of of several immune signaling proteins upon induction of KRAS^G12D^ including IL-6, IL-18, IL-23, IL-1B, IL-36β, and IL-36*γ*, which are involved in pro-inflammatory pathways, Treg differentiation and maintenance, and pro-angiogenic pathways [30, 36–39]. All of these signaling activities are associated with either tumor promotion or tumor suppressive effects. Therefore our model captures how cancer cells communicate with components of the immune microenvironment from the earliest stages of oncogenic transformation. While we have not directly tested these relationship in this study, our organoid model is amenable to co-culture with various other cell types, including components of the immune microenvironment, which can be leveraged to uncover the mechanistic details of these heterotypic cellular interactions.

## Materials and Methods

### Mice

Mouse husbandry and procedures were in accordance with protocols approved by the Unit for Laboratory Animal Medicine and Institutional Animal Care and Use Committee at the University of Michigan. Gt(ROSA)26Sor^tm4(ACTB-tdTomato,-EGFP)Luo^ (JAX#007676) and Kras^tm4Tyj^ (JAX#008179) mouse strains were obtained from the Jackson Laboratory and combined and maintained on a C57BL/6.

### Organoid isolation and Transduction

Primary mammary gland derived organoids were isolated in accordance with published protocols, with modifications [40]. Briefly, mammary glands #3, #4, and #5 were dissected from 8-13wk old mice. Lymph node was removed from gland #4 and all glands were pooled then minced ∼50 times with a #10 scalpel. Mammary tissue was then incubated in a 50-ml Falcon tube containing a collagenase solution for 1 h at 37°C while shaking at 180 RPM. Collagenase solution was composed of DMEM-F12 (10565-018; Gibco), 5% FBS (35-011-CV; Corning), 50 μg/ml gentamicin (15750-06; Glibco), 5 μg/ml insulin (I9278; Sigma-Aldrich), 0.1 g/mL collagenase (C2139; Sigma-Aldrich), and 0.1 g/mL trypsin (27250-018; Gibco). The solution was then spun down at 15000 rpm for 5 minutes. The fatty topmost layer was then transferred to a fresh collagenase solution for an additional 30 minutes at 37°C while shaking at 180 RPM. After this point, all tubes and pipette tips were coated in BSA (A6003-100G, Sigma-Aldrich) coat solution made in PBS. This second tube is then spun down at 1500 rpm for 5 minutes and pellets from the original tube and fatty layer are combined using DNase solution. DNase solution was composed of 2 U/mL DNase (04536282001, Roche) in DMEM-F12 (10565-018; Gibco). The suspension was incubated at room temperature for 3 min. Additional media was added to quench DNase activity and solution was centrifuged at 1500 rpm for 5 minutes . The pellet was resuspended in media, and epithelia was isolated through a series of 3-4 differential centrifugations (1500 × *g* for 10 seconds each). Organoid density was determined and adenovirus transduction was performed accordingly (20 μL virus (Ad-Cre: 1045; Vector Biolabs, Ad-Null: 1300; Vector Biolabs) plus 30 μL organoid growth media for every 800-1000 organoids). Organoid growth media contained DMEM-F12 (10565-018; Gibco), 1% penicillin– streptomycin (15140-122; Glibco), 1% insulin–transferrin–selenium–ethanolamine (51500-056; Gibco), and 2.5 nM FGF2 (F0291; Sigma-Aldrich). Transduction took place overnight in anti-adherence solution (354253; Corning) coated 96 well cell culture plates (229195; CELLTREAT). After transduction, organoids are collected and plated at desired concentration and extracellular matrix. Organoid media was supplemented with 2.5nM of FGF2 after an initial 24 hours in unsupplemented media. The media was changed on Day 4.

### 3D Organoid Culture

Primary mammary gland derived organoids were cultured following published protocols, with modifications. Briefly, growth factor reduced Matrigel (354230; Corning) was thawed in a quantity that allowed for plating of ∼100 organoids per 3D dome. Organoids were resuspended in 100uL of matrigel and plated on a 24 well tissue culture plate. Plates were plastic (229123, CELLTREAT) for culture only purposes and glass (662892; Greiner-One Bio) for imaging.

Matrigel domes were allowed to polymerize at 37°C for 30 minutes after which 500uL of organoid media was added per well. After 24 hours, media was changed to organoid culture media supplemented with 2.5 nM of FGF2 (F0291-25UG, Sigma-Aldrich). Organoids were cultured in this manner for up to 7 days.

### Immunofluorescence

Organoids in 3D Matrigel domes were fixed after 7 days in culture using 4% paraformaldehyde (PFA, 15714; Electron Microscopy Sciences) for 20 minutes shaking at ambient temperature.

Gels were then washed three times with PBS followed by permeabilization in 0.5% Triton X-100 (X100; Sigma-Aldrich) in PBS for 1 hour, gently shaking at ambient temperature. Samples were then blocked in 10% FBS (35-011-CV; Corning), 1% BSA (A6003-100G, Sigma-Aldrich), 0.1% Triton X-100 in PBS for 2-3 hours, gently shaking at room temperature. Primary antibodies were diluted in an antibody dilution buffer composed of 1% FBS, 1% BSA, 0.1% Triton X-100 in PBS and incubated at 4°C, shaking gently overnight. After three PBS washes, secondary antibodies were diluted in the antibody dilution buffer and incubated at ambient temperature, shaking gently, for 3-4 hours. The samples were then washed three times with PBS and stored at 4°C until imaging.

| Primary Antibody | Concentration |
|---|---|
| RAS G12D (MA536256; Invitrogen) | 1:100 |
| Phospho -Histone H3 (9701; Cell Signaling) | 1:400 |
| Cleaved Caspase-3 (966; Cell Signaling) | 1:400 |

Goat anti Rabbit 647 (A-21244; Thermo Scientific) was used as the secondary antibody and was diluted to a 1:500 concentration. F-Actin was stained with Phalloidin (PHDG1A, Cytoskeleton) and nuclei were stained with DAPI (#10236276001, Sigma).

### Confocal and Wide Field microscopy and quantification

Confocal images were acquired using a Yokogawa CSU-W1 spinning disk confocal microscope mounted on a Nikon Ti2 E Inverted microscope equipped with a LUN-F XL 405/488/561/640 laser combiner, Hamamatsu ORCA-FusionBT back-thinned camera, and Tokai-Hit full encolsed incubation system. Objectives used in the study include a LWD 20X CFI60 Plan Apochromat Lambda S and a LWD 40X water immersion CFI60 Plan Apochromat Lambda S . Images were acquired using the NIS Elements software. Individual organoid images were taken as z-stacks of 300μm and a z-step size of 20μm. Images of the entire gel were acquired with the CFI60 Plan Apochromat Lambda D 4X objective with a z-stack of 800μm and step size of 100μm.

Images were analyzed with Fiji software.

### Organoid size quantification

Maximum intensity z-stack projections of individual organoids were hand-outlined by a blinded individual for unbiased shape descriptor and area quantification. Since we were able to image the same organoid over time (day 1, day 3, day 5, day 7), the fold change in area was calculated using the day of interest area minus that same organoid’s area on Day 1. Fold change was normalized to account for variability in size between replicates and individuals by dividing fold change by that same organoid’s area on Day 1. Rate of growth was calculated using the normalized area on day n minus the normalized area of the same organoid on day n-2 divided by 2. For example, the first rate was the area of a given organoid on day 3 minus the area of that same organoid on day 1, divided by 2 to give the change in area per day.

### Fluorescence intensity quantification

Maximum intensity projections of individual organoid membranes (mT and mG) were used to segment organoids. Segmentation was performed using the Trainable Weka Segmentation (TWS) tool on Fiji. Organoid masks created from this segmentation were overlaid onto fluorescent channel original images and fluorescence intensity was measured only within these masks using the Fiji “Measure” tool. Raw integrated intensity was normalized to account for organoid size and depth in gel using the raw integrated density from the DAPI channel within the same mask.

### mG vs mT quantification

Maximum intensity z-stack projections of individual organoid mT channels and mG channels were segmented separately using TWS. Area of the segmented objects was measured using the Fiji “Measure” tool. Fluorescent area was normalized to account for organoid size and depth in gel using the same organoid’s Day 1 fluorescent intensity.

### Lumen formation quantification

Lumens were counted by going through single organoid z-stacks one z-slice at a time to determine if a lumen was present. To maintain consistency, a lumen was defined by an empty space enclosed completely by cells with at least one third of the perimeter being made of a single cell layer. Micro-lumens were defined as an empty space inside of the organoid enclosed completely by cells but had less than one third of its perimeter as a single cell layer.

### Statistical analysis

All statistical tests, except the ones performed for RNA-seq analysis, were performed using GraphPad’s Prism software. To test the normality of all samples, we performed a normality and lognormality test on all of the data. All data was non-parametric so Mann-Whitney tests were used to compare two data sets. The number of replicates (r) and number of measurements (n) are indicated in the figure legends. For lumen formation, chi-square tests were performed to determine if frequency of lumen occurrence was significant with Ad-Null condition as the expected number and Ad-Cre as the observed number. Significance stars for lumen formation are placed on the bar that represents the significantly *larger* population.

### RNA sequencing preparation and data analysis

Organoids were recovered from Matrigel by incubating them in cell recovery solution (354253, Corning) for 20 minutes. RNA from organoids was isolated using TRIzol Plus RNA purification Kit (12183555; Invitrogen) integrated with PureLink DNase treatment (12185010; Invitrogen). Purified RNA was quantified on a nanodrop, and sent to Plasmidsaurus for library prep and sequencing. Sequencing read quality was determined by FASTQC with default paramteres [41]. Sequencing adapters were trimmed using FASTP [42] also with default settings. Trimmed reads were aligned to the GRCm39 reference genome using RNA STAR (version 2.7.11a) in paired-end mode and default parameters with over 80% uniquely mapped reads for all conditions [43]. Aligned reads were quantified using featureCounts with the following parameters unstranded, paired-end and a minimum mapping quality per read of 10 [ 44]. Differential expression analysis was performed using DeSeq2 [45] with multi-factor design accounting for all experimental groups and experimental batches. Significant differentially expressed genes were defined using a p-value cutoff of <0.05 and log2FC of |1|. GSEA was used to determine pathway enrichment [46, 47], and PROGENy was used to determine intracellular signaling pathway involvement [23].

## Acknowledgements

We thank members of the West lab for their input into this research project. The West lab is supported by a Black in Cancer/Emerald foundation award.

---

**Supplemental Figure 1:**
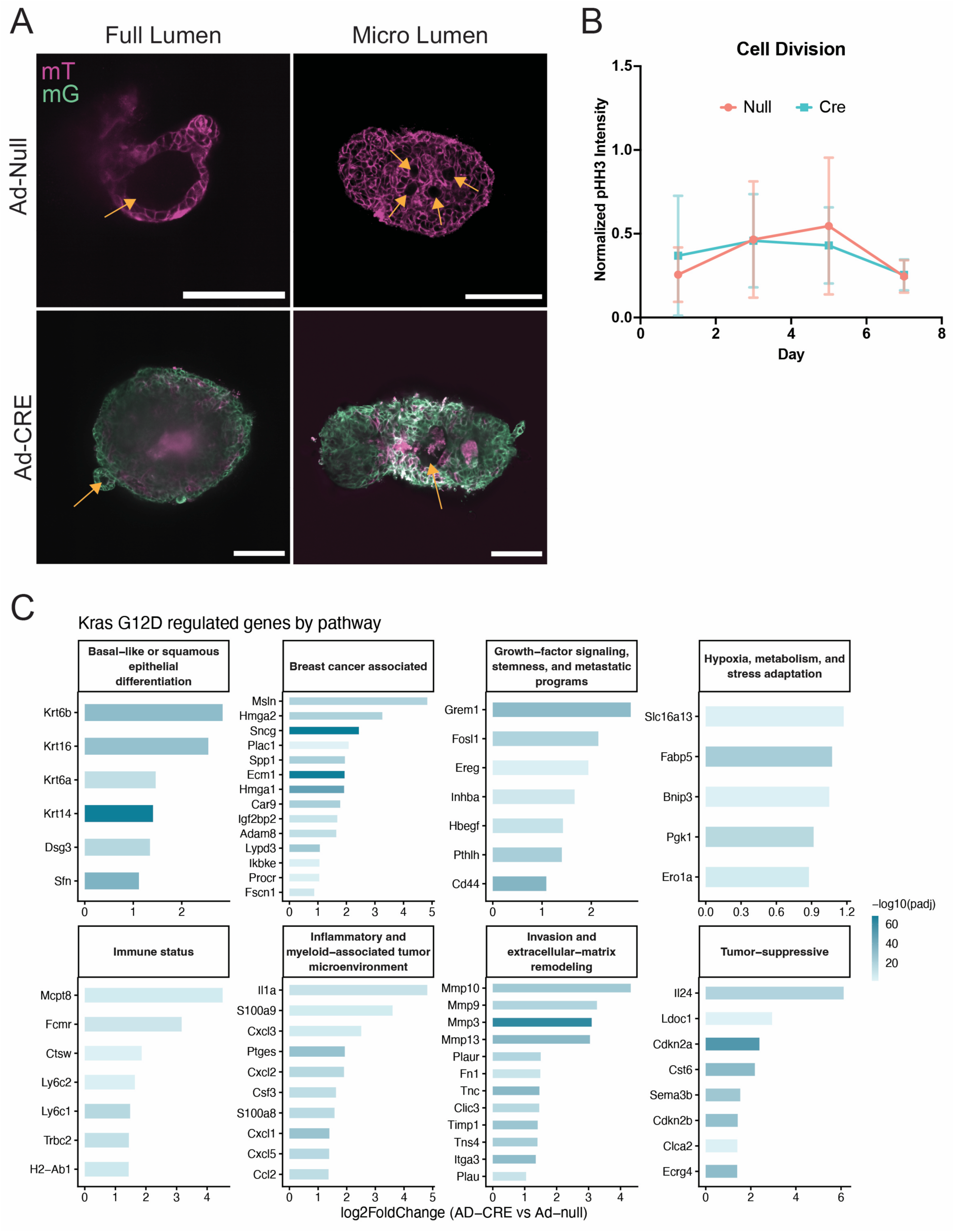
Morphological and molecular characteristics of KRASG12D expressing organoids. (A) Representative images of full lumens (left) and micro lumens (right), indicated by yellow arrows, from z-slices of organoids. Scale bar is 100 µm. (B) Quantification of phospho-histone H3 (pHH3) fluorescence over time. Error bars are SD. Mann-Whiteny test was used to determine that there was no significance difference in pHH3 fluorescence between Ad-Null and Ad-CRE organoids on any measured day. (C) Analysis of differentially expressed genes from a number of tumor-relavent biological pathways.

